# FloBuzz: A Modular Feeder System for Automated Aversive Conditioning in Bees

**DOI:** 10.64898/2026.02.08.704633

**Authors:** Elvin Gultekinoglu, Babur Erdem, Atakan Botasun, Okan Can Arslan, Sedat Sevin, Ayse Gul Gozen, Hande Alemdar, Erol Sahin, Ali Emre Turgut

**Affiliations:** Department of Mechanical Engineering, Middle East Technical University, Ankara, Türkiye; Center for Robotics and Artificial Intelligence (ROMER), Middle East Technical University, Ankara, Türkiye; Department of Pharmacology and Toxicology, Faculty of Veterinary Medicine, Ankara University, Ankara, Türkiye; Department of Biological Sciences, Middle East Technical University, Ankara, Türkiye; Department of Computer Engineering, Middle East Technical University, Ankara, Türkiye

**Keywords:** Artificial flower, Honey bee, Electric shock avoidance, Computer vision

## Abstract

Understanding how environmental stressors affect pollinator behavior is essential for assessing ecosystem health. Automated flower and robotic feeder systems (AFRFSs) have transformed how pollinator foraging and learning are studied. Still, most systems remain reward-centric, limiting their ability to probe aversive learning and nociception under field conditions. Here, we present FloBuzz, a modular AFRFS that couples automated reward delivery with computer-vision-based visit detection to trigger closed-loop electric-shock stimulation in free-flying honey bees. In our setup, FloBuzz consisted of a 3D-printed feeder with a shock grid, a syringe pump with fluid-level feedback, and an electric shock stimulus trigger module that applied user-defined shock patterns. In a proof-of-concept trial that alternated between shock-free, 6 V, and shock-free, 9 V intervals, bee visitation increased over time during shock-free periods but declined during shock periods, with a steeper decline at 9 V than at 6 V, demonstrating a voltage-dependent avoidance. By enabling programmable, time-resolved aversive stimulation at an artificial flower in outdoor conditions, FloBuzz expanded AFRFS capabilities beyond purely reward-based paradigms. Evidently, FloBuzz’s modular design will permit the investigation of diverse behavioral paradigms, including studies of toxin exposure, cognitive plasticity, and reward processing.

## 1. Introduction

Pollination is one of the most critical factors for plant survival, and bees, in particular, are considered keystone species for ecosystem sustainability and agricultural productivity (Aizen et al., 2009). Therefore, studying bee behavior is necessary for both environmental sustainability and agriculture. Automated flowers and robotic feeder systems (AFRFSs) are essential tools for improving experimental conditions for pollinators (Chapman et al., 2023). Here, we introduce FloBuzz, an AFRFS capable of investigating aversive conditioning in bees. Aversive conditioning is a method to examine an organism’s ability to learn to avoid a negative stimulus (Abramson, 1986). In our study, FloBuzz enabled these aversive conditioning studies to be conducted directly in outdoor environments.

AFRFSs are essential experimental tools that enable researchers to study the mechanisms underlying pollinator behavior, learning, and floral trait evolution under controlled, replicable conditions (Chapman et al., 2023; Essenberg, 2015). Compared to natural flowers, AFRFSs offer superior control over reward timings and floral cues such as color, shape, scent, and nectar concentration, thereby mitigating challenges in foraging experiment design. This level of control makes AFRFSs invaluable for dissecting how pollinators integrate multimodal cues during decision-making and for quantifying foraging efficiency across sensory modalities (Broadhead & Raguso, 2021; Ômura & Honda, 2005; Raguso & Willis, 2002). Additionally, working with real flowers poses substantial logistical challenges due to correlations between floral traits (Frey & Bukoski, 2014), high maintenance costs (Chapman et al., 2023), and numerous uncontrolled environmental variables (Nordström et al., 2017). Natural variations in nectar chemistry, volatile release, or petal morphology can bias behavioral interpretations (Silva & Dean, 2000). Therefore, AFRFSs provide consistency across experimental replicates and eliminate this bias. Moreover, AFRFSs can repeatedly present controlled combinations of visual, olfactory, and reward cues in response to animal visits (Debeuckelaere et al., 2024; Essenberg, 2015). In previous studies, the utility of AFRFSs has been expanded for behavioral ecology and neuroethology by enabling long-term experiments (Giurfa & Núñez, 1992), real-time monitoring (Sokolowski & Abramson, 2010), and standardized reward delivery (Giray et al., 2015).

Furthermore, foraging decisions in natural environments are shaped not only by rewards but also by risks, highlighting the need for AFRFSs that can integrate aversive or nociceptive stimuli alongside controlled rewards. Integrating nociceptive stimuli and predator dummies into AFRFSs enables the investigation of aversive learning. For example, imitating sit-and-wait predators such as mantids or crab spiders as aversive elements suppresses honey bee foraging (Bray et al., 2014; Huey et al., 2017; Ings & Chittka, 2008). Other methods apply nociceptive stimuli, such as heating (Foley et al., 2025; Gibbons et al., 2022).

Electric shock is another nociceptive stimulus in the electric shock avoidance assay, which is a standard laboratory procedure. It has been utilized to investigate the role of chemicals (Erdem et al., 2026; Nouvian et al., 2018) and neuromodulators (Agarwal et al., 2011), caste specificity (Dinges et al., 2013; Avalos et al., 2017), and neural pathways (Plath et al., 2017) on learning. Furthermore, the electric-shock avoidance assay has recently been adapted for free-flying bumblebees in indoor settings (Sarlak et al., 2023).

We devised FloBuzz to adapt the standard laboratory electric shock avoidance assay for use in outdoor environments. Through its computer-vision-based visit detection and trigger module, the system delivers shocks in user-defined continuous, intermittent, or delayed patterns. FloBuzz also provides food solutions as appetitive stimuli alongside these aversive triggers. Thus, FloBuzz integrates various engineering approaches to combine the well-established method of electric shock avoidance with appetitive stimuli through an automated system, enabling researchers to conduct studies in natural environments. Consequently, FloBuzz is bound to facilitate research on learning, foraging, predatory incidence, and risk assessment in a natural environment for free-flying honey bees.

## 2. Method

### 2.1. System Architecture of FloBuzz

FloBuzz was designed to support controlled behavioral experiments by integrating stimulus (electric shock) delivery via computer-vision-based triggering, and automated food supply as a reward into a single modular system (Figure 1). The system consisted of a feeder module, the In-Trigger module, and a syringe pump with a control unit (Figure 2). The feeder module was the primary component that enabled interaction with bees. An electric shock grid in the feeder module was connected to a shock switch, which is controlled by the In-Trigger module. The feeder module was attached to the syringe pump via a flexible food delivery tube.

**Figure 1.**
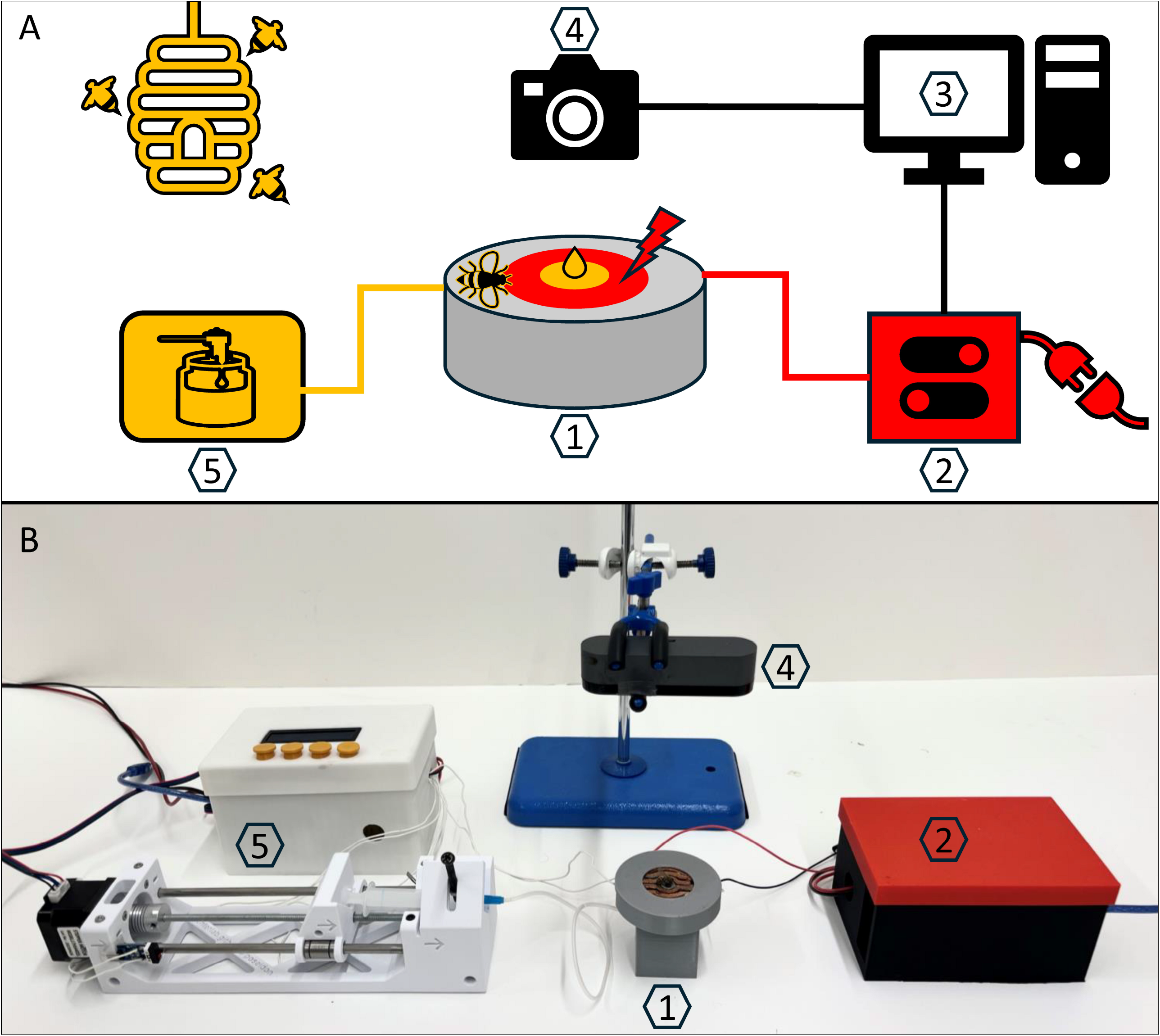
View of the experimental setup. The schematic illustration of the experimental setup (A) and the picture of the experimental setup (B). The feeder module provides the food solution to the bees and acts as the platform where the electric shock is applied (1). The shock switch triggers an electric shock based on control signals from the In-Trigger module (2). The computer runs the In-Trigger module and processes visual input from the webcam (3). The webcam acquires visual information from the feeder module for visit detection (4). The syringe pump supplies the food solution to the feeder module, regulated by a control unit (5).

**Figure 2.**
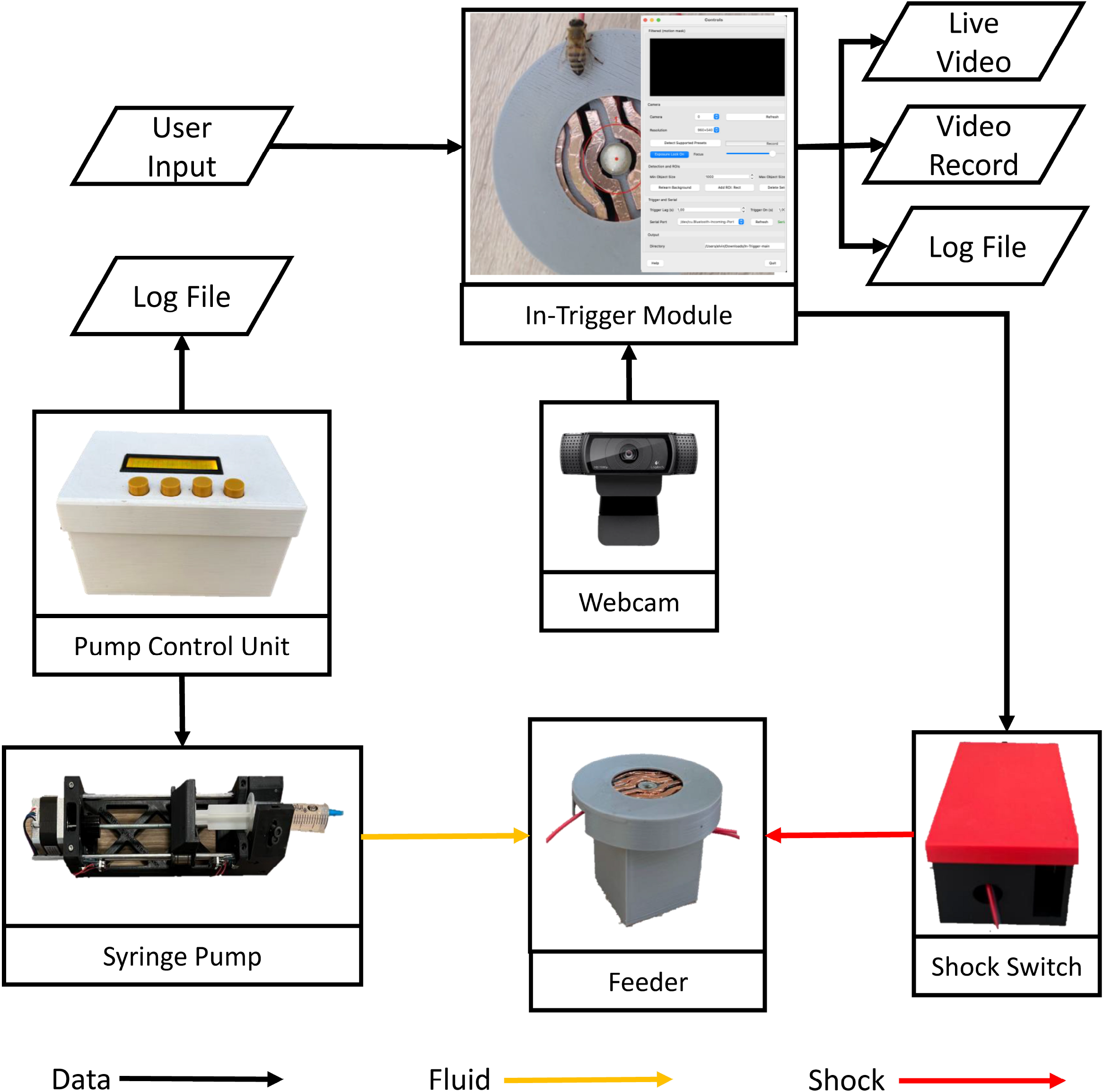
Low-level representation of modules and their interactions in the FloBuzz system. The feeder module is connected to the syringe pump and the shock switch. The shock switch is controlled by the In-Trigger module and enables the activation and deactivation of the electric stimulus. Visual information from the feeder module is acquired by a webcam and transmitted to the In-Trigger module. The pump control unit governs the operation of the syringe pump, which supplies the food solution to the feeder module continuously.

The In-Trigger module detected objects entering predefined regions of interest (ROI) within the camera frame using computer vision algorithms. It activated the shock switch with user-defined delays and durations.

The syringe pump delivered a steady supply of food solution to the feeder module. Its design was adapted from the open-source Poseidon syringe pump (Booeshaghi et al., 2019). A pump control unit (described in the Supplement File 1) enabled the pump to automatically refill the feeder as bees consume the solution. A script running on this subsystem monitored solution levels and logged each refilling event.

### 2.2. Feeder Module

The feeder module served as the physical interaction interface between the bees and the FloBuzz system, providing both the food reward and the aversive electric stimulus. The feeder module was composed of four parts that fit together: a main base, a cap, an electric shock grid, and a collecting box (Figure 3).

**Figure 3.**
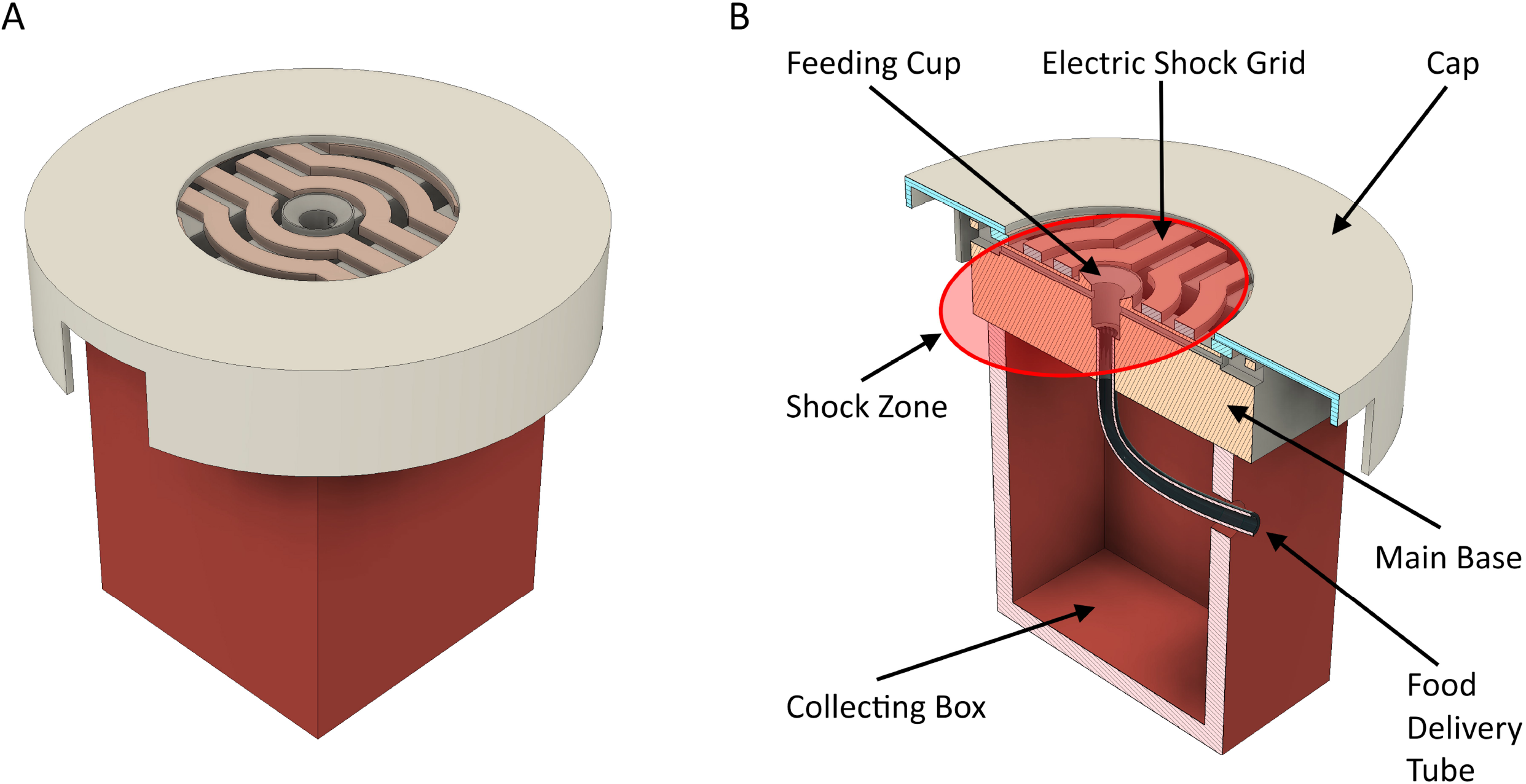
Technical drawing of the feeder module (A) and the sectional view (B). The main base includes the feeding cup. The food delivery tube is penetrated to the main base and it enables controlled food solution delivery. The electric shock grid surrounds the feeding cup and displays the shock zone. The cap is positioned above the grid and it defines the landing area. The collecting box beneath the assembly provides a large internal cavity to capture any overflow of the food solution.

In the module, the main base housed the feeding cup and included holes for silver-plated feedback cables. These cables function as electrodes and provide analog readings to an Arduino board mounted on the custom PCB (Supplementary Figure S1). The feedback cable lines are also interfaced with the pump control unit to enable closed-loop actuation of the syringe pump during automatic refilling. The feedback algorithm monitored the food solution and triggered automatic refilling when the cup was empty by evaluating analog readings against an experimentally determined threshold value. The feeding cup was positioned at the center of the main base and functioned as the reservoir for the food solution. The feeding cup geometry featured a gentle slope and edge protrusions to minimize surface tension and prevent overflow. The feeding cup was 4.5 mm in diameter and 5.3 mm deep; these dimensions prevented bees from falling in while limiting evaporation. Feedback cables were placed 1.2 mm below the rim of the feeding cup, allowing bees to comfortably consume the solution while maintaining continuous monitoring of the solution level. An opening at the bottom of the feeding cup secured a butterfly-type infusion tube, hereafter referred to as the food delivery tube, to enable controlled delivery. The food delivery tube transported the food solution from the syringe pump to the feeding cup of the feeder module.

The electric shock grid delivered the aversive stimulus with a 2.5 mm gap between grid lines. The bees received a shock when their feet completed the circuit by bridging two separate lines. The electric shock grid can be manufactured by applying copper tape to a 3D-printed grid or by directly cutting a copper plate. The cables were soldered directly to the grid and connected to the shock switch (Supplementary Figure S2).

The FloBuzz feeder cap enclosed the main base. It featured a circular design to ensure the shock area is radially symmetric. This geometry was chosen because bees preferentially visit radially symmetric artificial flowers (Wignall et al., 2006; Møller & Sorci, 1998). With a diameter of 68.0 mm, the cap provided ample landing space for the bees. The hole in the center of the cap was 33.0 mm in diameter. This hole exposed the electrical grid, and its radius was approximately the length of a bee, ensuring that the bee would touch the grid in this shock zone. A catch basin beneath the main base captured any overflow, preventing food solution from spilling into the experimental area.

### 2.3. Computer Vision-Based Stimuli Trigger Module (In-Trigger)

To enable precise, automated aversive stimulus delivery, we developed In-Trigger, a system integrating a video-processing pipeline, a graphical interface, and microcontroller-based actuator control (available from https://github.com/baburerdem/In-Trigger). This integrated module offers a flexible, low-latency, and fully configurable system for closed-loop experimental paradigms that require precise synchronization between behavioral detection and stimulus delivery. The system detects objects entering predefined spatial regions within the camera frame. It activates external stimulus devices with user-defined delays and durations (Figure S3).

We implemented the core functionality in Python using the OpenCV (Bradski, 2000) library for real-time video acquisition and analysis. The script could manage background subtraction, object detection, Region of Interest (ROI) mapping, trigger logic for the shock switch, and data logging. Video was captured from USB cameras via the highest-priority available backend. Motion detection was performed using a *MOG2* (Zivkovic & van der Heijden, 2006) background subtraction model. Detected foreground masks were morphologically filtered and intersected with user-defined ROIs. The system tracked individual moving objects by matching detected contours across consecutive frames. It evaluated their position relative to each ROI. A trigger event was confirmed only when a detected object was localized within a specific ROI.

Each ROI was mapped to a unique digital output pin on an Arduino controller, which receives commands via serial communication and switches the stimulus device accordingly. When an ROI trigger was confirmed, a shock event was scheduled on the Arduino controller through a sequence of activation and deactivation commands. Timing parameters, including the lag between detection and trigger onset, and the duration of the active trigger, were configurable via the GUI.

The GUI developed by *PySide6* enabled full control of experimental parameters and system status. Users can select cameras and resolutions, define and edit ROIs interactively on the live preview, toggle horizontal flipping, and adjust autofocus or manual focus settings. Controls also included configuration for detection thresholds (minimum and maximum object size), trigger delays, stimulus durations, and the selection of a serial port for Arduino communication. The GUI provided real-time visual feedback of both raw and processed frames, displaying annotated bounding boxes and ROI outlines (Figure 4).

**Figure 4.**
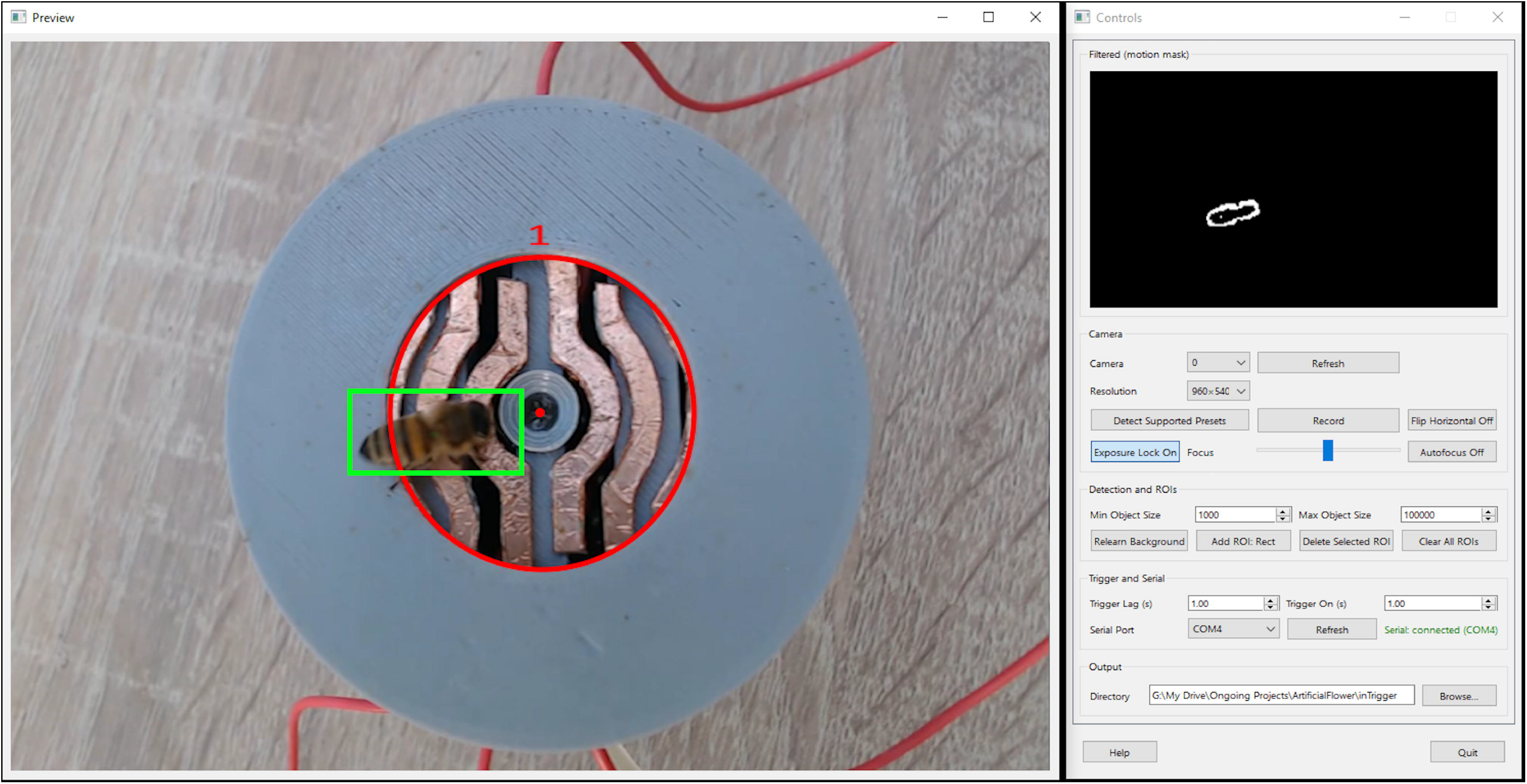
In-Trigger graphical user interface and real-time detection during the experiment. The left panel shows the live camera preview of the feeder module during an experimental session. The circular red region indicates the predefined shock zone. A visiting honey bee is detected and is enclosed by a green bounding box. The right panel displays the In-Trigger control interface, which provides real-time visualization of the filtered motion mask (top) and allows full configuration of experimental parameters, including camera selection and resolution, background relearning, minimum and maximum object size thresholds, trigger lag and on-time durations, serial communication with the shock switch, and data recording options.

All relevant experimental data were automatically logged during recording sessions. Video output was saved in MP4 format, alongside a high-quality JPEG snapshot of the first annotated frame to verify ROI placement. Concurrently, a timestamped plain-text log file was created, detailing frame-by-frame object counts per ROI, shock activation status, frame indices, and the associated video file. These logs ensured traceability and facilitated post-hoc validation of each experimental run.

The electric shock switch was controlled by an Arduino board, which sends on/off commands to the IRF520 driver. An external power source and a MOSFET IRF520 driver module are generally used as the solid-state switching element. Gate-drive and protective components were implemented, and the switching system was controlled through the In-Trigger GUI (Figure S2).

### 2.4. Field Experiment

To validate our system, we conducted a field experiment in a small apiary at the Middle East Technical University campus. The feeder module, syringe pump, and a camera were located on a table near the hives (Figure 1). The syringe pump was initiated, and the feeder module maintained a 50% (v/v) honey/water solution. Bees were allowed to approach the artificial flower freely. Based on preliminary observations of the duration between landing and feeding, the shock lag-time and on-time were set to 4 and 3 seconds, respectively. The In-Trigger module successfully detected entries into the shock zone (Supplementary Video S1). When an electric shock was applied, bees dispersed from the feeder in response to the aversive stimulus. Simultaneously, the In-Trigger module recorded the sessions for subsequent analysis.

### 2.5. Statistical analysis

All statistical analyses were conducted in RStudio using the R version 4.3 (R Core Team, 2020). Normality was evaluated using the Shapiro–Wilk test. Correlations were quantified using the Pearson correlation test for normal data (*p* ≥ 0.05) and the Spearman correlation test when normality assumptions were not met (*p* < 0.05).

### 2.6. Ethical Note

According to Article 4.b of the Regulation on the Working Procedures and Principles of Animal Experimentation Ethics Committees, published in the Official Gazette of the Republic of Turkey (February 15, 2014, Issue: 28914), an experimental animal is defined as “any non-human vertebrate organism, including free-living or reproducing larval forms, live cephalopods, and mammals from the last third of their normal fetal development, used in procedures.” Therefore, research conducted with honey bees (*Apis mellifera*) is not subject to evaluation by the Ethics Committee. We also declared that no bee individuals were harmed in our study, and the experiment durations were kept short enough to obtain only the necessary data, minimizing the discomfort experienced by the bees during the experiment.

## 3. Results

We manually scored the 450-second video recording of the electric shock-avoidance experiment at 5-second intervals. The number of bees on the flower and whether the electric shock was active were noted. The experiment included consecutive intervals as follows: a 75-second shock-free interval, then a 165-second 6 V shock-applied interval until the bees were no longer on the flower, a second 135-second shock-free interval during which the bees were expected to return to the flower, and a 60-second 9 V shock-applied interval during which the bees escaped (Figure 5). To quantify visitation trends, correlation coefficients were calculated between the number of bees on the flower and time for each experimental interval.

**Figure 5.**
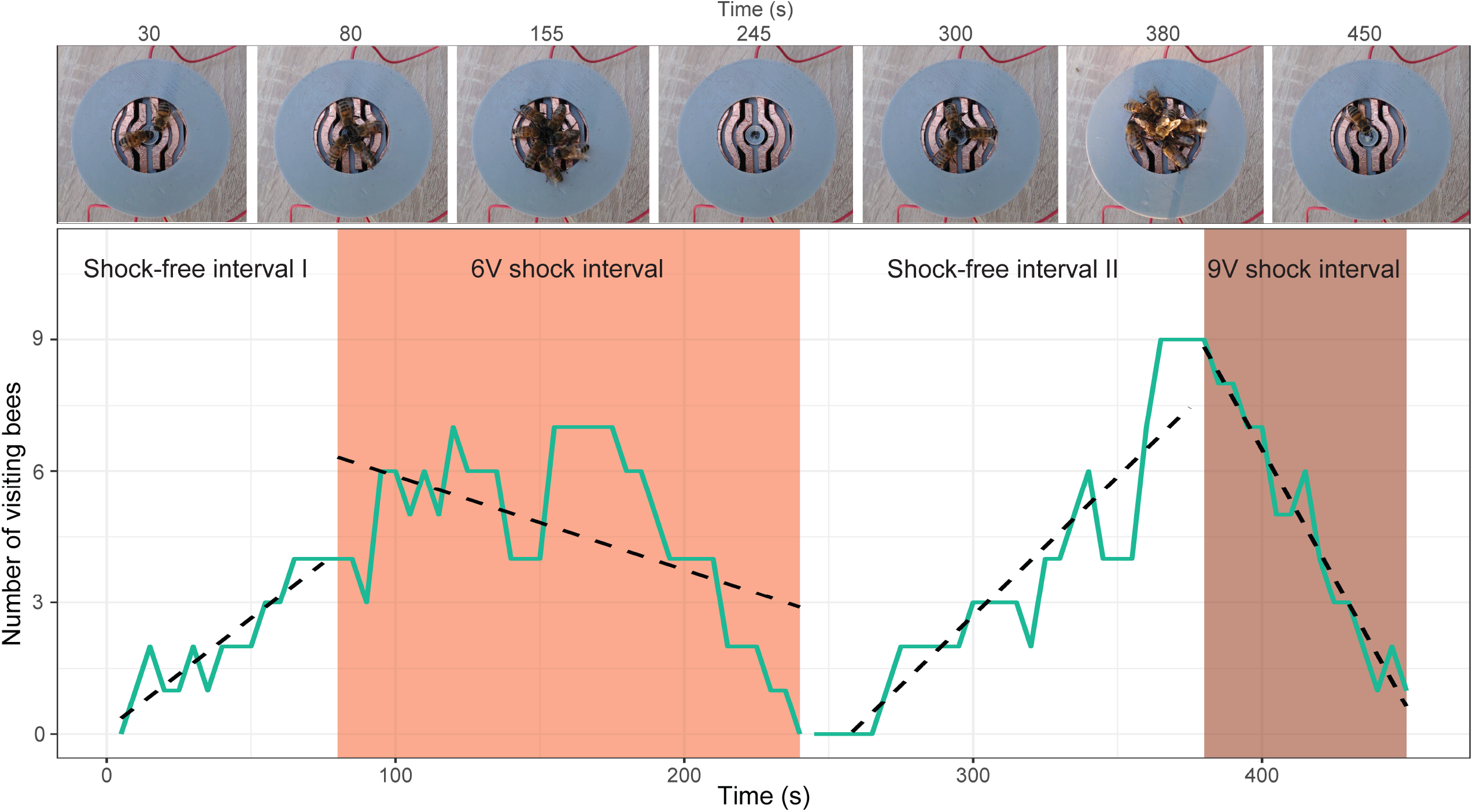
Visitation curve of the bees to the feeder module. The green line shows the change in the number of visiting bees over time. Red shaded areas indicate shock-applied intervals. Dashed black lines are correlation lines.

A positive correlation by time was found in the first shock-free interval (Pearson *r*(13) = 0.910, *p* < 0.0001) and the second shock-free interval (Spearman *ρ*(25) = 0.961, *p* < 0.0001). Because the correlation coefficients were very close (0.910 and 0.961), it can be concluded that the patterns of bee visits to flowers are quite similar. A negative correlation was found in the 6 V shock-applied interval (Spearman *ρ* (31) = −0.437, *p* = 0.008) and the 9 V shock-applied interval (Pearson *r*(13) = −0.976, *p* < 0.0001). The higher correlation coefficient at 9 V (0.976) than at 6 V (0.437) indicated that the higher voltage elicited a stronger avoidance response.

The proof-of-concept trial showed that the system elicits clear, voltage-dependent changes in visitation. Bee visits increased over time during shock-free intervals but declined when the feeder delivered an electric shock. This decline was sharper at 9 V of electric shock than at 6 V. That was consistent with prior studies reporting that the shocking action supports robust aversive learning and increasing aversive intensity amplifies avoidance and shifts decision thresholds (Abramson, 1986; Dinges et al., 2013; Gibbons et al., 2022; Kirkerud et al., 2017; Plath et al., 2017).

## 4. Discussion and Conclusion

We designed FloBuzz as a modular, field-deployable AFRFS to enable controlled bee–flower interaction experiments under natural conditions, coupling precisely timed, programmable aversive stimulation with reward delivery. The integration of a standardized feeder module, an electronically switchable shock grid, and a vision-triggered control pipeline (In-Trigger) allows the system to operationalize closed-loop stimulus delivery. The electric shock stimulus is activated contingent on the bee’s presence at the feeder, rather than on a fixed schedule. Our design addresses a significant practical bottleneck in behavioral ecology by empowering behavioral assays on free-flying bees that were previously restricted to laboratory arenas.

Compared with the broader AFRFS literature, FloBuzz’s differentiator is that it is not primarily a reward-centric, cue-manipulation, or remotely operable AFRFS; instead, it adds a programmable nociceptive channel and implements contingent (visit-triggered) aversive stimulation under field conditions. Many AFRFS and artificial-flower approaches are optimized for standardized reward delivery, manipulation of multimodal floral cues, long-term repeated trials, and monitoring of visitation dynamics (Essenberg, 2015; Chapman et al., 2023; Debeuckelaere et al., 2024; Sokolowski & Abramson, 2010; Ohashi et al., 2010). The advantage of FloBuzz is its explicit targeting of aversive paradigms in fields conventionally confined to laboratory settings (Abramson, 1986; Agarwal et al., 2011; Dinges et al., 2013; Kirkerud et al., 2013, 2017; Plath et al., 2017). Evidently, it complements field-associated aversive approaches based on predator cues or heat-based nociception (Ings & Chittka, 2008; Bray & Nieh, 2014; Huey & Nieh, 2017; Gibbons et al., 2022; Foley et al., 2025).

The FloBuzz system was tested in the field and revealed clear, voltage-dependent avoidance at the feeder. Bee visitation increased during shock-free intervals and declined during shock application, with the 9 V stimulus eliciting a significantly more pronounced reduction than the 6 V stimulus. This proof-of-concept demonstrates that FloBuzz can deliver reliable, time-resolved aversive stimulation to free-flying bees in an outdoor setting while maintaining automated reward delivery. The observed pattern is consistent with established evidence that increased aversive intensity strengthens avoidance responses and alters decision thresholds in shock-based assays (Abramson, 1986; Dinges et al., 2013; Plath et al., 2017; Gibbons et al., 2022).

A key advantage of FloBuzz is its combination of modular hardware that can be adapted to diverse feeder geometries and closed-loop triggering driven by real-time computer vision rather than manual operation. Furthermore, the In-Trigger module is species-agnostic and can be integrated into experimental setups for a wide variety of animals.

These features enable users to perform trials that are difficult to implement with reward-centric systems alone, including controlled aversive–reward trade-off assays. This approach lays the groundwork for future research on how pollinators balance risks, costs, and rewards under realistic environmental conditions, and how stressors such as agrochemicals and predation risk reshape fundamental trade-offs.

## Authorship Contribution Statement

A.B.: Writing – original draft. Investigation. A.E.T.: Writing – original draft. Methodology, Conceptualization, Resources. A.G.G.: Writing – original draft. B.E.: Writing – original draft. Visualization, Software, Methodology, Investigation, Formal analysis, Conceptualization. E.G.: Writing – original draft. Visualization, Software, Hardware, Methodology, Investigation, Conceptualization. E.S.: Writing – original draft, Methodology. H.A.: Writing – original draft, Methodology. O.C.A.: Writing – original draft, Investigation. S. S.: Investigation.

## Acknowledgements

This work was supported by the Middle East Technical University Research Fund [grant number: ADEP-302-2024-11468]; the EU grant RoboRoyale [grant number: 964492]. We would like to thank our intern Khuslen Tugsjargal for his help.

## Data availability

Script of the In-Trigger module, printable 3D models of the parts used in the FloBuzz feeder, experimental data, and statistical analysis script are available in the Zenodo repository: https://doi.org/10.5281/zenodo.18458890. Also, the In-Trigger scripts that will include possible future updates can be accessed from the GitHub repository: https://github.com/baburerdem/In-Trigger

