## Supplementary File for "FloBuzz: A Modular Feeder System for Automated Aversive Conditioning in Bees"

**Supplements**

**Syringe Pump and Pump Control Unit**

The food solution level in the feeding cup is measured by silver-plated multi-stranded copper cables that are electrodes attached to the cup. A 1 KΩ resistor was placed in series with the cable connected to the analog pin of the microcontroller (Arduino NANO - ATmega328P) to limit excess current when the bee bridges the sensing electrodes through the solution (Figure S1).

The control unit continuously samples the feeding cup through a feedback cable connected to an analog pin, and compares it to an empirically selected threshold (~1000, 10-bit ADC). When the signal crosses the threshold indicating depletion, the firmware enables the stepper to refill. Once the feeding cup is full, the stepper motor is stopped, and the pump is turned off to prevent overfilling. Refilling is performed in a pulsed manner: the pump is driven for a fixed on-time and then paused for a fixed off-time. The duty-cycled operation prevents overshoot and allows the analog readings to settle.

The control unit is integrated into a custom PCB, which consists of several key components: a microcontroller-based platform, a power regulation stage, a stepper motor driver (A4988), user interface components, and external input ports. The PCB uses an LM2575S-5V switching regulator to provide a stable 5V DC output for logic components. The stepper motor driver controls the precise movement of the syringe plunger, while the user interface includes four push buttons and a 2×16 I2C LCD for displaying operational data. The user interface allows for manual intervention in addition to automatic pump start. In emergencies, the user can operate the pump in either pull or push mode (the Arduino script is available in the Zenodo Repository at <https://doi.org/10.5281/zenodo.18458890> ).

The system’s operational log is collected and stored through serial communication between the firmware and a Python script (Given in the Zenodo Repository at <https://doi.org/10.5281/zenodo.18458890> ). The system log includes the operational state (on/off), the active mode, and the time duration of each mode synchronized with global time. To enable data logging, the Arduino on the PCB must be connected to a computer through its USB interface, allowing serial communication between the firmware and the Python script. Serial communication was configured at 9600 baud to exchange logs between the firmware and the host PC.

All components of the syringe pump were fabricated by Creality K1 Max 3D printer with PLA filament, enabling precise geometries, lightweight construction, and cost-effective prototyping. For practical use, the syringe needle is removed and replaced with the tubing from a butterfly infusion set, which simplifies handling and allows the fluid line to be replaced frequently whenever necessary.

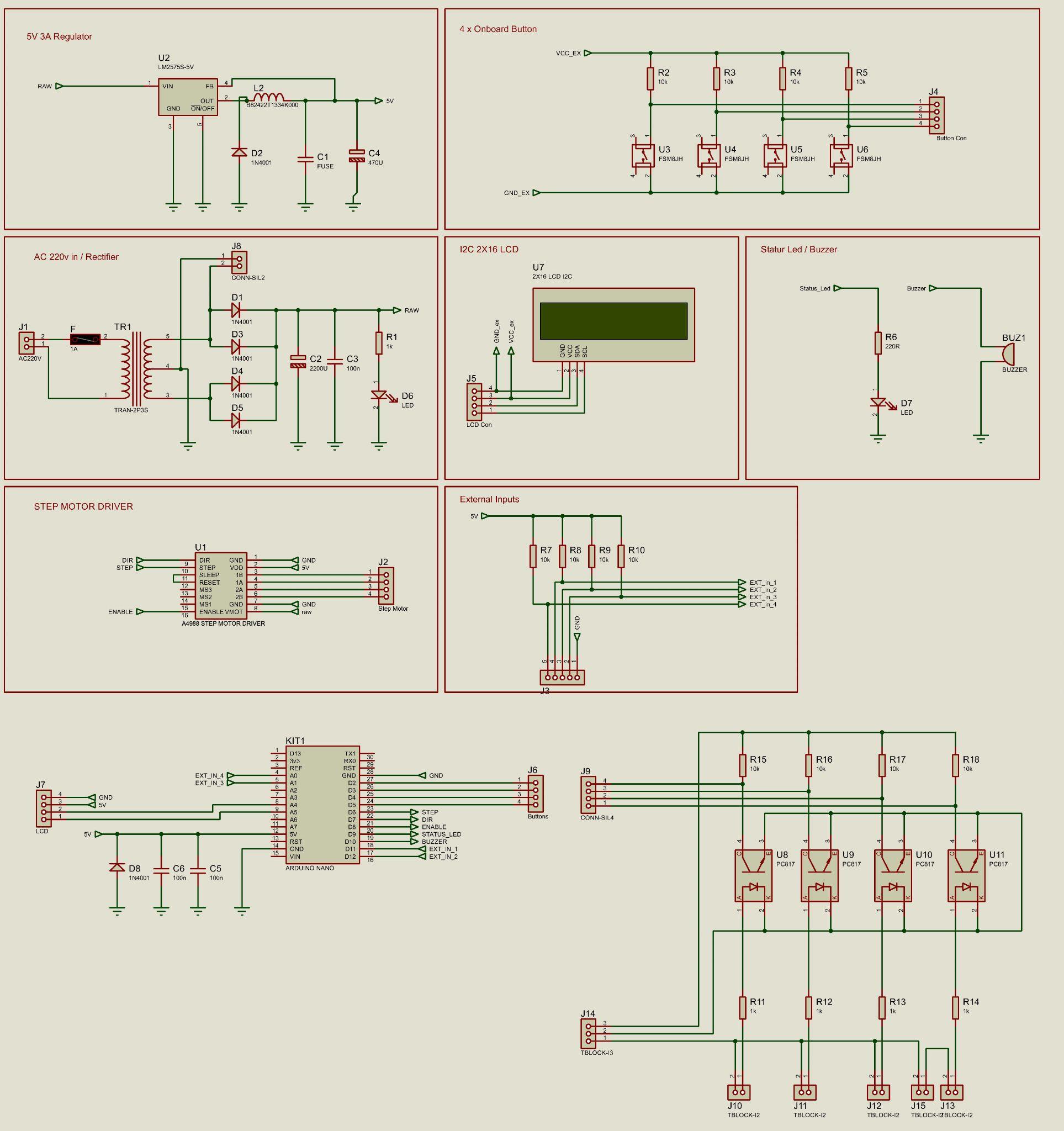

Figure S1. Custom PCB design of syringe pump control unit.

**In-Trigger Module**

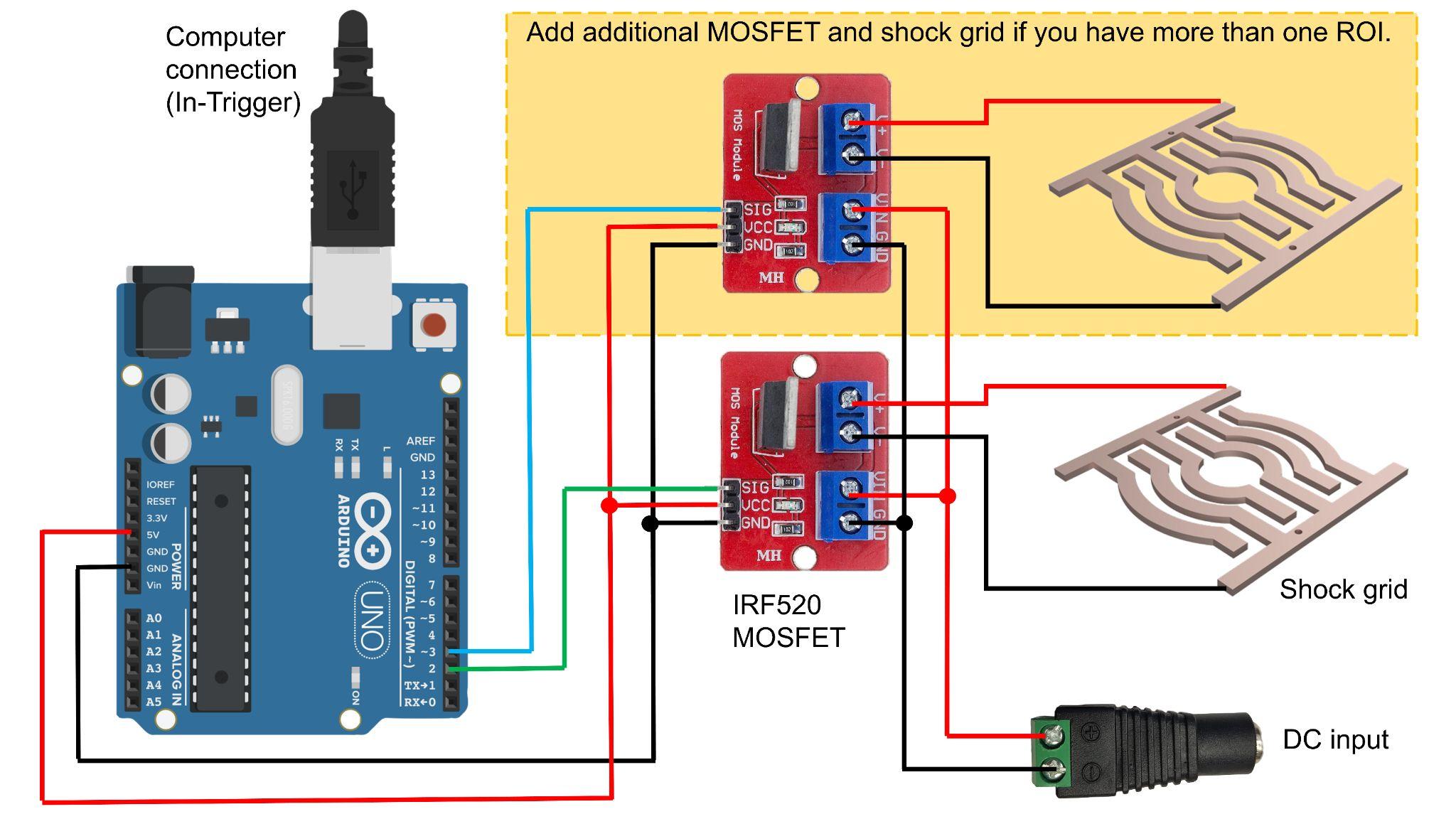

Figure S2. Circuit diagram of the shock switch module.

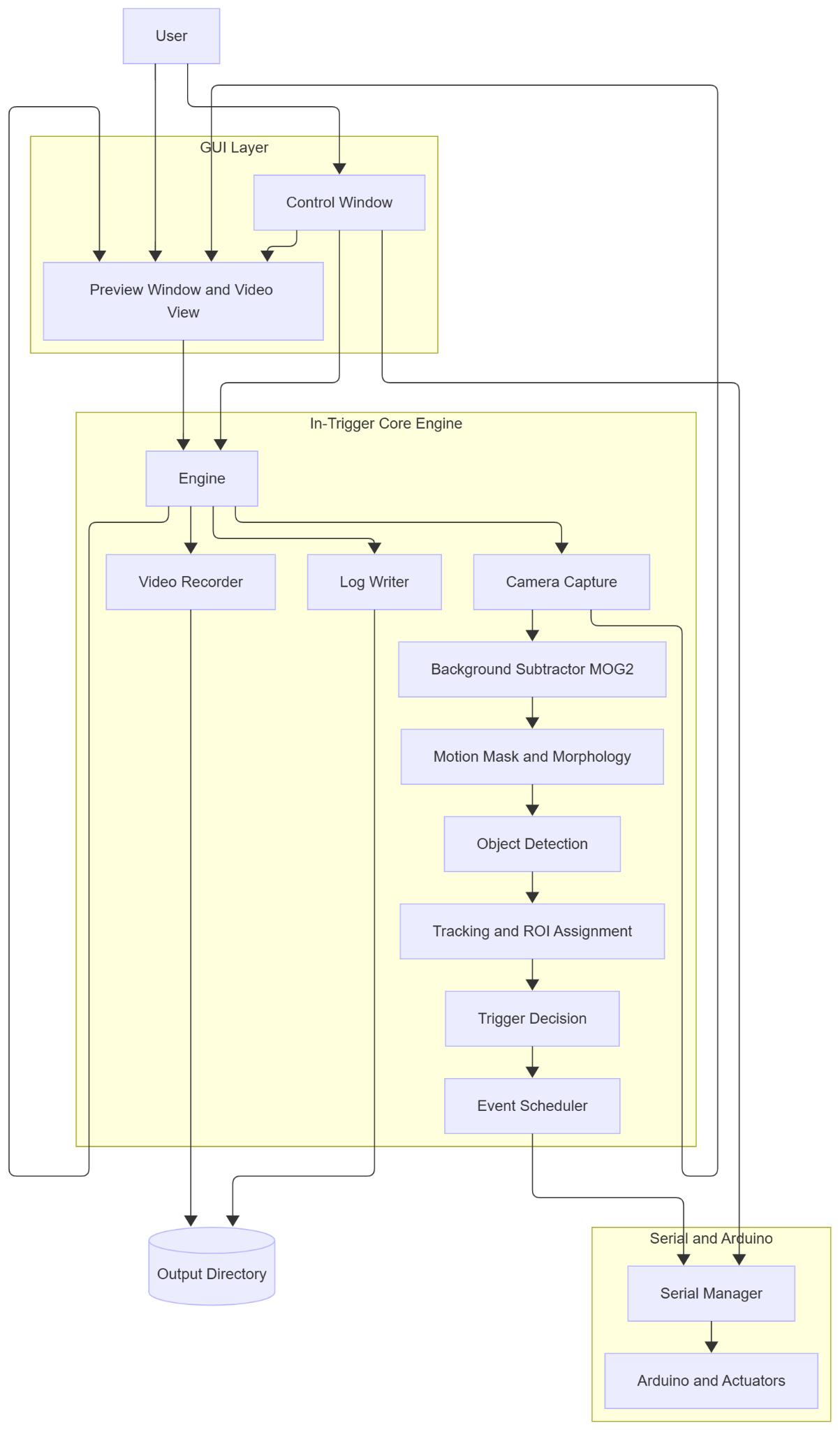

Figure S3. Flowchart of the In-Trigger module.
